# Envenomation leads to venom protein reduction and recovery delay in bumblebee workers

**DOI:** 10.1101/2025.07.25.666811

**Authors:** Safira Moog, Mario Dejung, Amitkumar Fulzele, James Carolan, Thomas J. Colgan

**Affiliations:** Institute of Organismic and Molecular Evolution, Johannes Gutenberg University, Hanns-Dieter-Hüsch-Weg 15, 55128 Mainz, Germany; Institute of Molecular Biology (IMB), 55128 Mainz, Germany; Department of Biology, Maynooth University, Co Kildare, Maynooth, Ireland

**Keywords:** venom, bumblebee, envenomation, proteomics

## Abstract

The evolution and deployment of venom systems in animals has been central to their ecological success, though not without costs. For social bees, venom plays a key role in nest defence yet for species, such as honeybees, the act of stinging generally leads to death. However, for other social bee species, such as bumblebees, where stinging is not apparently fatal, it raises the question of what, if any, are the costs associated with venom usage (envenomation). Here, we investigated proteomic changes in the venom sac, the primary reservoir for venom storage, of *Bombus terrestris* workers over a short time-course post-envenomation. Our analysis reveals three key findings: 1) envenomation leads to changes in the venom sac proteome with affected proteins generally reduced in the bumblebee venom sac, suggestive of a molecular cost; 2) quantified reductions of traditional venom-associated proteins post-envenomation indicate changes in venom composition post-usage; and 3) recovery of reduced venom-associated proteins post-envenomation was delayed further indicating a potential cost at the molecular level associated with usage. Overall, our findings provide new insights into the consequences of venom usage in bumblebees.

## 1. Introduction

Predation and competition place selective pressures on individuals that lead to adaptive strategies and traits to enhance survival [1]. One such trait is venom, an evolutionary innovation, which has evolved in numerous organisms to aid in predation, as well as in defence, acting as a strong deterrent against predators and nuisances through inducing pain, paralysis, and even death [2,3]. Venoms are highly complex mixtures of bioactive compounds containing a high number of small peptides, proteins, and toxins, as well as polyamines, organic molecules, and salts [4]. Venomous animals actively inject their self-synthesised venom into targets through the process of envenomation, disrupting normal physiological or biochemical processes [4], serving key roles in predation and defence. The evolutionary and ecological importance of venom is underlined by its independent evolution at least 101 times across diverse animal lineages [3].

Although venom use provides clear advantages in predation and defence, maintaining and using the venom system is costly, both in terms of product synthesis, as well as post-use regeneration. For example, increases in metabolic rate have been described in snakes, spiders, and scorpions during venom regeneration [5–7], potentially taking several days, such as 16 days in the spider *Cupiennius salei* [8] and up to 28 days in snakes [9]. This suggests that venom usage is energetically costly and, therefore, should not be used wastefully. Venomous animals have evolved strategies favouring conservative venom use through physiological, morphological, and behavioural adaptations, to minimise costs, as postulated in the venom optimisation hypothesis [10]. An increasingly recognised concept supporting the venom optimisation hypothesis is that venom use can be regulated depending on the ecological context [3,9]. The state of the envenomating organism and that of the prey are both suggested to play a role in venom use as venom regulation can be affected by prey size and prey type. For example, the spider *C. salei* alters the amount of venom it injects depending on the size of the prey likely to minimise venom use [10]. Furthermore, from an evolutionary perspective, several secondary losses or reductions of morphological traits associated with venom synthesis or delivery have occurred, such as in a marbled sea snake caused by a dietary switch [11], implying that venom is energetically costly to maintain at an evolutionary scale [9]. While energetic expenditure by predators could be compensated by the consumption of prey [9], there is no such direct compensation in terms of reward for venom usage during defence. Therefore, animals using defensive venoms may be more likely limited in evolving mechanisms for venom regulation as failure of defence leads to potential death [3], suggesting that venom usage in this context may be more costly.

An ecologically and biomedically relevant group that employ venom usage in defence are bees (Clade Anthophila), a taxon-rich group (20,000 described species [12]), of important ecological and commercial pollinators. While venom usage by bees is predicted to have evolved initially for hygienic purposes, such as in the sterilisation of nesting sites [13,14], social bee venoms are generally associated with defence causing pain, tissue damage, and potentially death [15]. Like other hymenopterans, the venom system of bees evolved from an ancestral ovipositor and accessory sex glands. Thus, the venom system is only present in females while males lack the apparatus and are unable to sting [16,17]. Despite the importance of venom, venom usage in bees has largely been studied from a biomedical perspective, with our knowledge of venom use in an ecological context limited. The best studied bee species, honeybee *Apis mellifera*, highlights the cost placed on individuals in terms of self-sacrifice during nest defence. In honeybees, morphological adaptations and associated defence tactics have evolved to maximise pain in vertebrate threats, leading to a high individual cost in the form of fatality after stinging through autonomy (self-amputation) of the stinging apparatus [18]. However, such morphological adaptations are not present in other stinging social bees, such as bumblebees, which do not possess the same modified barb found in honeybee workers which results in the ability to repeatedly sting both invertebrate and vertebrate threats [19,20]. Compared to other social bees, such as honeybees, which may number in the thousands, bumblebee colonies are considerably smaller (∼150-300 workers [21]), which means workers may be of greater individual importance to colony productivity. However, our understanding of potential costs associated with venom usage within bumblebee workers is currently unknown. Such costs may be reflected in changes in venom composition placing energetic demands for regeneration, as well as limiting reuse or potency.

To understand whether envenomation results in changes in bumblebee venom composition, as well as venom regeneration time, we employed mass spectrometry-based proteomics to investigate changes in the venom sac proteome of *Bombus terrestris* workers over a short time-course (1h, 24h, 168h) after envenomation. We chose the venom sac as it acts as a reservoir for the storage of venom proteins that are primarily produced in the connected tubular venom glands [22]. Therefore, protein changes in the venom sac proteome can provide an insight into the molecular response and recovery post-envenomation. As venom regeneration is potentially costly [9,10,23], we predict that envenomation leads to: 1) changes in the proteome composition of the venom sac, including reduced abundances of known venom proteins; and 2) a delay in the recovery of venom protein abundances to pre-envenomation levels due to energetic costs.

## 2. Material and methods

### (a) Colony maintenance

*Bombus terrestris* colonies (n=5) were obtained from Koppert Biological Systems (Netherlands) and kept under constant conditions at 28°C and under red light illumination. Sugar water (APlinvert, Germany) and pollen (Biobest, Belgium) were provided *ad libitum*. To control for age, we collected callows (within 48h post-eclosion) and transferred each to a box (15cm x 15cm x 15cm) consisting of four related mature workers (distinguishable due to clipped wings), brood, and sugar water for approximately 72h to allow for maturation [24].

### (b) Envenomation procedure

To induce envenomation, we used parafilm-covered 0.2ml tubes filled with 100µl autoclaved phosphate buffered saline (PBS) to collect venom from each worker. For each collection, we temporarily restrained the bee and placed the collection tube close to the aculeus. We visually confirmed stinging by the presence of a hole in the parafilm. For control bees, they experienced the same handling procedure, but no venom was collected. Post-envenomation, workers were returned to their boxes and remained until a designated sampling time-point, which was either 1h, 24h or 168h after procedure (one week; Fig.1). At each designated collection time-point, workers were transferred into individual cryotubes, snap-frozen in liquid nitrogen, and stored at -80°C before being dissected. In total, we collected 25 workers (Envenomation group: 1h (n=5), 24h (n=4) and 168h (one week, n=3); Control group: 1h (n=5), 24h (n=4) and 168h (n=4)). Individuals, one each from a different colony, were used for each treatment group and time-point. We also assessed the collected venom for protein content using the Qubit protein assay kit.

**Fig. 1.**
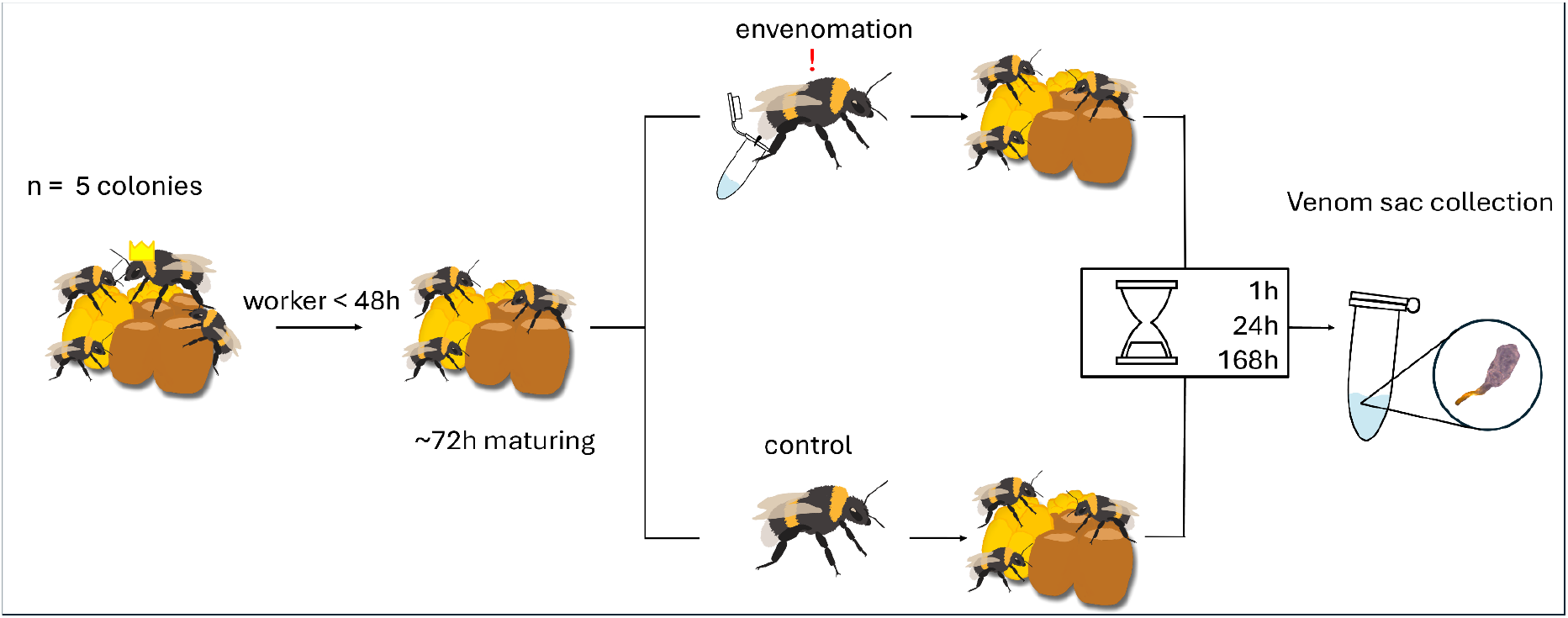
Envenomation procedure and venom sac collection. Worker callows (<48h after eclosion) of five different colonies were collected and kept for 72h under colony like conditions until the envenomation and control treatment was performed. In the envenomation treatment, a bee stung into a parafilm covered tube while control bees did not sting. The bees were returned to a nursery colony and sampled at 1h, 24h and 168h, post-treatment. The venom sac was later dissected and analysed using mass-spectrometry-based proteomics.

### (c) Venom sac extraction and proteomic analysis

For each bee, the venom sac was extracted and subsequently homogenised in PBS. To determine differently abundant proteins, an EASY-nLC 1000 system (Thermo Scientific) and a Q-Exactive Plus Orbitrap mass spectrometer (Thermo Scientific) were used. MS/MS data were analysed using MaxQuant (v.1.6.10.43; [25]; Full details of sample preparation and analysis are provided in Supplemental Material). In total, across all samples, the venom sac proteome contained 2,031 proteins, including 36 previously described *B. terrestris* venom proteins [26] (Supplemental Material, Supplemental Table S1).

### (d) Statistical analysis and data visualisation

To identify differentially abundant proteins between treatments at each time-point, we performed Welch’s t-tests comparing the imputed LFQ values of envenomation and control bees in R. P-values were corrected for multiple testing using a false discovery rate of 0.05. For data visualisation, we used the R packages ggplot2 (v.3.5.1; [27], ggpubr (v.0.6.0; [28]) and ggrepel (v.0.9.6; [29]).

### (e) Functional annotation of described venom sac proteins

To determine biological processes associated with venom sac proteins, we performed a Gene Ontology (GO) term enrichment analysis using a Fisher’s exact test (Benjamini-Hochberg adjusted-*p* < 0.05) implemented using the “weight01” algorithm of the topGO R package (v.2.54.0; [30]). We performed individual GO term enrichment analysis for genes coding for proteins with elevated or reduced abundances between treatments for each time-point, respectively, using scripts from [31].

## 3. Results

To understand how the venom sac proteome changes at individual time-points (1, 24, and 168h post-envenomation), we performed pairwise comparisons of treatment and control bees at each time-point finding that envenomation collectively resulted in moderate changes in the venom sac proteome with between 5-8% of identified proteins changing across the three time-points examined. We identified significantly differentially abundant proteins (SDAPs) (pairwise t-test, FDR<0.05) at each time-point (1h:153 SDAPs; 24h:103 SDAPs; 168h:71 SDAPs) with a significantly higher number of SDAPs (76-79% of all SDAPs) having reduced abundance in the venom sac post-envenomation at all time-points (Binomial exact tests: *p*<0.05; 1h:116 SDAPs, 24h:81 SDAPs, 168h:56 SDAPs) suggestive of depletion of specific proteins or reduced translational activity. To understand the biological processes that such proteins are involved in, we performed GO term enrichment analyses finding that genes coding for proteins reduced after 1h and 24h were significantly enriched for “cytoplasmic translation” (GO:0002181) (Fisher’s exact test, BH-adjusted *p*<0.05; Supplemental Table S1). After 168h post-envenomation, there was no significant enrichment of GO terms for reduced SDAPs (Fisher’s exact test, BH-adjusted *p*>0.05). Despite general patterns of reduced protein composition in the venom sac post-envenomation, we also found proteins that increased in abundance after venom usage (1h:37 SDAPs, 24h:22 SDAPs, 168h:15 SDAPs). There was no significant enrichment of biological process GO terms for elevated proteins at all time points (Fisher’s exact test, BH-adjusted *p*>0.05). Nearly all reduced and elevated SDAPs were unique to individual time-points with few exceptions (Supplemental Figure 2).

As our analyses revealed general reductions in protein abundance post-envenomation, which may be expected due to venom use and depletion, we examined abundance changes of previously described venom proteins. Prominent components of bumblebee venom, such as phospholipase A2-like (XP_012170988.1) at 1h and bombolitin/melittin (XP_020718386.1) at 24h post-envenomation, were significantly reduced (pairwise t-test, BH-adjusted<0.05) within the venom sac. We also found other proteins, previously described in bumblebee venom, such as venom carboxylesterase (XP_020724240.1), platelet derived growth factor (XP_003394009.1), and protein yellow (XP_020719667.2), which were reduced at 1h or 24h post-envenomation before correction for multiple testing (pairwise t-test, BH-adjusted *p*>0.05; Fig.2A-B), but were not significantly different after correction. In contrast, no differences in expression levels for these proteins were observed at 168h post-envenomation between control and treatment bees (pairwise t-test, BH-adjusted *p*>0.05; Fig.2C). Lastly, one venom protein increased in abundance post-envenomation, a trehalase-like protein (XP_003400853.1), which was elevated 24h post-envenomation (Fig.2B).

**Fig. 2.**
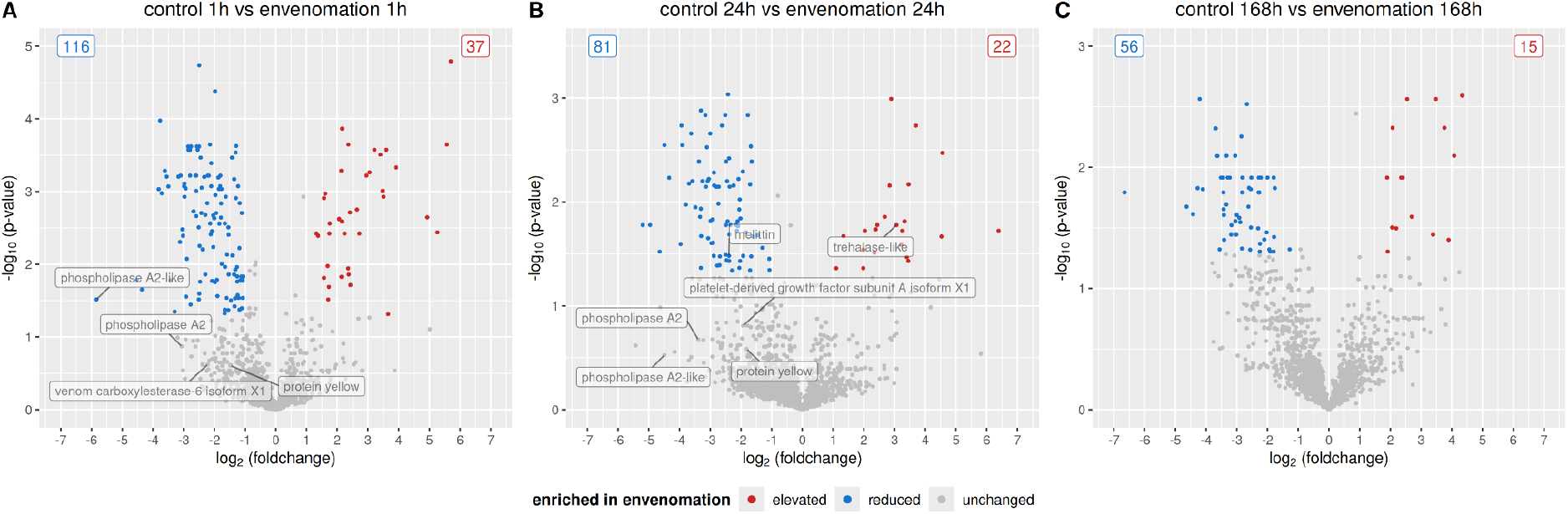
Envenomation is associated with relatively larger reductions in protein abundance in the venom sac of *Bombus terrestris* workers at three different time-points post-treatment. **(A)** Volcano plot showing the protein abundance differences (log_2_ fold change) between workers post-envenomation relative to control workers 1h, **(B)** 24h, and **(C)** 168h post-envenomation. Proteins with significantly (pairwise t-test, FDR<0.05) reduced and elevated abundance post-envenomation are individually coloured (blue = reduced abundance; red = elevated abundance; grey = unchanged). Previously described venom proteins in *B. terrestris* [26], which were differentially abundant before correction for multiple testing, are individually labelled.

## 4. Discussion

Many animals across different lineages use venom, mainly for predation and defence, highlighting the ecological and evolutionary importance of this biological innovation. Nevertheless, synthesis of venom and maintaining a venom system is energetically costly [9,10,23]. Given this energetic requirement [9,10,23], the aim of this study was to examine the potential consequences associated with venom usage (envenomation) in *B. terrestris* workers, where venom is mainly used for individual and colony defence. Therefore, we performed a proteomic-based analysis to understand how the proteome composition of the venom sac, the primary reservoir of venom proteins, changes for *B. terrestris* workers after envenomation, but also to determine whether envenomation leads to short-term changes that may affect reuse. We found that envenomation results in changes in the venom sac proteome with relatively larger reductions in protein abundances, including prominent components of bumblebee venom, suggestive of potential costs at least at the molecular level. Our analysis also highlights a delay in recovery of venom abundances as evident by reduced abundances after 24 hours post-envenomation, which are recovered at least one week later indicating a delay in venom regeneration, which may have consequences for short-term potency and reuse.

During venom delivery, venomous organisms contract muscles of the venom reservoir to allow for the regulated release of venom to structures that facilitate delivery [3]. In our analysis, of the proteins that changed after envenomation, the majority were reduced, including notable reductions of previously described venom components, which is in line with the possible depletion of stored venom proteins due to venom usage. Among the venom proteins reduced in the venom sac within 24 hours, we identified phospholipase A2 and bombolitin, which are among the most abundant proteins in bumblebee venom [32]. Phospholipase A2 induces inflammatory responses [33] with bumblebee phospholipase A2 exhibiting apoptotic activity in mammals [34]. Melittin, the honeybee’s homologue of bombolitin, is the most abundant peptide in honeybee venom and is known to induce strong and painful reactions in target organisms [33] which, given the close relationship, could also be the case for bombolitin. In addition, bombolitin enhances phospholipase A2 activity [35] suggesting an important role of both components in venom potency.

The observed short-term reduction of venom proteins after envenomation may result in less potent venom which may not induce as much pain and be less toxic in their targets. For example, snakes may use dry bites to conserve venom usage [36] or deliver lower dosages during venom regeneration [5]. The availability of less venom could lead to behavioural changes of the individual. Although not examined in the present study, there is evidence that some venomous animals are aware of their venom status resulting in distinct behaviours if less venom is available, such as in some snakes, which hunt smaller prey or use constriction after they used venom or when venom is limited [3]. In bumblebees, it could be that individuals are less likely or unable to appropriately defend the nest directly post-envenomation. However, this individual cost in repeated defence may be offset or compensated for by nestmates, which can act cooperatively to deter colony-level attacks. Further research is required in this aspect to understand cooperative group work in nest defences with respect to how it influences venom usage and regeneration.

Venom usage is often associated with regeneration time after use. While our analysis found reductions at two points within 24 hours post-envenomation, venom proteins were observed to have similar abundance levels with control individuals after seven days post-envenomation. While our analysis did not include intermediate time-points, meaning that recovery may be sooner, our results indicate that recovery of key venom components occurs within a week of envenomation. As it takes time to replenish the venom composition, there are periods when venom cannot be used or when less venom may be available. Timing of venom regeneration varies across species with regeneration potentially taking up to several weeks [8,9]. Speed of regeneration is also influenced by metabolic rate. For example, the metabolic rate of the scorpion *Parabuthus transvaalicus* increases by 21% - 40% in the first days of venom regeneration [6,7], highlighting metabolic costs to replenish venom. While we did not directly measure metabolic activity, proteins involved in translation were differentially abundant immediately after envenomation, which may reflect investment of energetic resources to replenish used venom. However, as bumblebees are social insects, the recovery of venom proteins may not only be affected by metabolic costs but also by their social environment. For example, in the social acorn ant *Temnothorax longispinosus*, venom production was associated with social cues. Venom investment by an individual was linked to social parasite pressure whereby colonies that experienced a social parasite raid had more estimated venom per individual. In addition, venom investment after a raid was associated with the task that a single individual performs with foragers having more venom than nurses [37]. Thus, potential threats and social factors, such as a low number of individuals, may influence venom production and venom recovery of an individual to ensure the colony’s viability. Bumblebees typically use venom to defend their nests, and the colony usually consists of numerous individuals that are able to contribute to the collective nest defence which could enable the investment in venom, including the time required for replenishment, to be distributed among nestmates. In our study, workers were kept in small social groups which may lead to a faster venom recovery to be capable of defending in the event of an attack.

During venom regeneration, energetic costs could be compensated by increased food consumption. At 24h post-envenomation, the venom protein trehalase was elevated in the venom sac. Trehalase is an enzyme also found in the haemolymph of insects which hydrolyses trehalose, the primary storage sugar in insects found in the haemolymph and in venom glands [38], into glucose and is associated with the hunger state and appetite regulation in honeybees [39–41]. Low trehalose levels correspond with an increase in octopamine levels in the honeybee brain leading to elevated appetite levels [39–41]. Thus, elevated trehalase levels in response to envenomation could lead to a higher availability of glucose, which is the main energy resource in insects, and simultaneously to lower trehalose levels resulting in an enhanced appetite to compensate energetic costs for venom regeneration. As bumblebees use venom for defence and do not offset energetic costs by nutritional gain from successful prey capture and consumption, this could be a mechanism to ensure offsetting energetic costs. Higher food consumption can offset the negative energetic effects of immune stimulation in *B. terrestris* [42,43] and, therefore, may potentially represent a mechanism to compensate for the induction of metabolically costly activities. In our experiments, small micro-colonies with low worker density were used that were provided with ample food, which could expedite recovery in at least one week, whereby in the natural environment, under restricted resources, recovery time may take longer. Future studies would benefit from examining how venom regeneration is affected by resource limitation.

## CONCLUSIONS

Venom plays an essential role in the defence of social insects, such as bumblebees, yet usage can result in consequences observable at different biological levels. Using proteomic-based profiling of an organ fundamental in the storage and usage of venom, we show that envenomation and subsequent regeneration is associated with changes in protein abundance, with the majority being reduced, including known venom proteins. Our findings also indicate a potential delay in venom recovery, which may have implications for repeated use as well as the potency of bumblebee venom during regeneration. Future work will benefit from direct examination of venom products, including bioassays to determine shifts in potency after envenomation as well as examine how venom usage may affect other life history traits, such as longevity or reproduction, which is of particular interest as the reproductive system in bees shares a close evolutionary history with that of venom. Collectively, our findings provide a novel insight into the proteomic composition of an organ essential for venom delivery, highlight how venom synthesis and storage changes post-envenomation, as well as indicate a potential delay in venom regeneration, which likely has implications for the capacity of bumblebees to defend themselves and their nests.

## Ethics

This work did not require ethical approval from a human subject or animal welfare committee.

## Data accessibility

Raw mass spectrometry data will be made available via the PRIDE database (PXD066016). Additional methods, results, data, and analysis code are available in the electronic supplementary material.

## Authors’ contributions

S.M.: conceptualization, data curation, investigation, methodology, formal analysis, project administration, visualization, writing—original draft, writing—review and editing; M.D.: data curation, methodology, formal analysis; A.F.; data curation, methodology; J.C.: methodology, formal analysis; T.J.C.: conceptualization, formal analysis, funding acquisition, investigation, methodology, project administration, resources, supervision, writing—original draft, writing—review and editing.

## Acknowledgements

We thank the IMB Proteomics Core Facility for proteomics analysis. Funding of the German Research Foundation supported the Q Exactive Plus system (DFG Project #240874965). We thank Jasmin Cartano for handling sample submission, as well as Jiaxuan Chen for feedback on an earlier draft of the manuscript. We also thank Ina Köhler for the use of the bumblebee designs.

