## Supplemental materials for "Envenomation leads to venom protein reduction and recovery delay in bumblebee workers"

### Preparation of samples for proteomic analysis

Protein quantities of venom sac samples were measured and 5µg protein per sample were resolved briefly in a 4-12% NuPAGE Bis-Tris gradient gel, stained with Coomassie blue and cut into small gel cubes, followed by destaining in 50% ethanol/25mM ammonium bicarbonate. Afterwards, proteins were reduced in 10mM DTT at 56°C and alkylated by 50mM iodoacetamide in the dark at room temperature. Enzymatic digestion of proteins was performed using trypsin in 50mM ammonium bicarbonate at 37°C overnight. Following peptide extraction sequentially using 30% and 100% acetonitrile, the sample volume was reduced in a centrifugal evaporator to remove residual acetonitrile. The peptide solution was purified by solid phase extraction in $C_{18}$ StageTips.

Peptides were separated on an EASY-nLC 1000 system (Thermo Scientific) via an in-house packed 30-cm analytical column (inner diameter: 75µm; ReproSil-Pur 120 $C_{18}$-AQ 1.9-µm beads, Dr. Maisch GmbH) by online reverse phase chromatography through a 105-min non-linear gradient of 1.6-32% acetonitrile with 0.1% formic acid at a nanoflow rate of 225nl/min. The eluted peptides were sprayed directly by electrospray ionization into a Q Exactive Plus Orbitrap mass spectrometer (Thermo Scientific). Mass spectrometry was conducted in data-dependent acquisition mode using a top10 method with one full scan (scan range: 300 to 1,650m/z; resolution: 70,000, target value: $3 \times 10^6$, maximum injection time: 20ms) followed by 10 fragment scans via higher energy collision dissociation (HCD; normalized collision energy: 25%, resolution: 17,500, target value: $1 \times 10^5$, maximum injection time: 120ms, isolation window: 1.8m/z). Precursor ions of unassigned or +1 charge state were rejected. Additionally, precursor ions already isolated for fragmentation were dynamically excluded for 20s.

MS/MS data were analysed using MaxQuant (v.1.6.10.43; [1]) with match between runs activated. Mass spectra were searched against a target-decoy database containing the forward and reverse sequences of *B. terrestris* proteome predicted from the genome assembly (Bter_1.0; [2]) obtained from Ensembl Metazoa release 53, as well as a default list of contaminants. As certain venom-associated proteins, such as bombolitin, have truncated gene models within the *B. terrestris* genome assembly, we included proteins from two melittin homologues in *Bombus impatiens*. MaxLFQ algorithm was employed for protein quantification, using its default normalisation option and requiring a minimum LFQ ratio count of 1. Detected proteins were filtered for reverse binders and contaminants. MaxQuant classified a protein as expressed when at least one unique peptide for each protein was detected or when a single peptide was identified across multiple samples by multiple MS/MS spectra in the same matching group (six matching groups in total: one group per treatment and time-point). Missing values were imputed in two route steps. Proteins not detected in a single replicate were imputed with a normal distribution at mean of the measured replicates. If a protein was not detected in more than one sample, then the values were imputed close to the detection limit.

### Comparison with previous venom proteomes

As known venom-associated proteins would be expected to be detected in our venom sac analysis, we compared a list of identified proteins in our dataset with venom-associated proteins previously described for *B. terrestris* [3]. To ensure high confidence in previously identified venom-associated proteins, we downloaded data from van Vaerenbergh et al. (2015) [3] and filtered identified proteins based on high peptide-spectrum matches (PSM value >= 20), resulting in the retention of 43 high-confidence venom-associated proteins (Supplemental Table S1). We then examined the presence of these proteins in our venom sac proteome dataset to identify their presence, as well as examine if their abundance changed post-envenomation.

### Principal component analysis

To examine differences in proteome profiles between treatments and across time-points, we first performed a principal component analysis (PCA) using the imputed LFQ intensity values of all quantified proteins. In addition, we generated Euler diagrams to examine overlap between proteins with significantly elevated and reduced abundance in response to envenomation for each time-point generated in R using the Eulerr package (v.7.0.2; [4]).

### Gene Ontology term enrichment analysis

To determine biological processes influenced by the act of envenomation, we performed Gene Ontology (GO) term enrichment analyses using Fisher's exact tests implemented in R using the package topGO (v.2.46.0; [5]) with the "weight01" algorithm. Due to the depauperate nature of the annotations assigned to non-model organisms, such as bumblebees, we downloaded GO terms assigned to genes in the fruit fly *D. melanogaster* and then transferred them to their homologues in the *B. terrestris* genome Bter_1.0 [2]. Genes with no annotated GO term or genes coding for proteins that lack homology with proteins of *D. melanogaster* were excluded. We then used this generated database to test for enrichment of GO terms for genes of interest. In total, we performed GO term enrichment analyses for: 1) all genes coding for identified protein groups found in the venom sac (protein group list of the MaxQuant output) to determine biological processes associated with proteins found in the venom sac; 2) and proteins that had elevated or reduced abundance post-envenomation across each time-point. All GO enrichment analyses were conducted using a node-size of 50.

### PCA result

We examined global changes in the venom sac proteome in workers post-envenomation by performing a principal component analysis using all 2,031 proteins identified across all treatments (control and envenomation) and time-points (1h, 24h and 168h post-treatment). Our analysis revealed proteome-wide expression changes with samples clustering by time-point, with the first principal component (PC1), which explained 28.05% of the variance, separating the latest time-point (168h post-treatment) from the others, indicating temporal changes in the venom sac. Furthermore, the second principal component (PC2) largely separated individuals from the later envenomation treatments (24h and 168h post-treatment; Fig. S1) providing the first indication that envenomation results in changes in the proteome composition, which are prolonged after the act of envenomation.

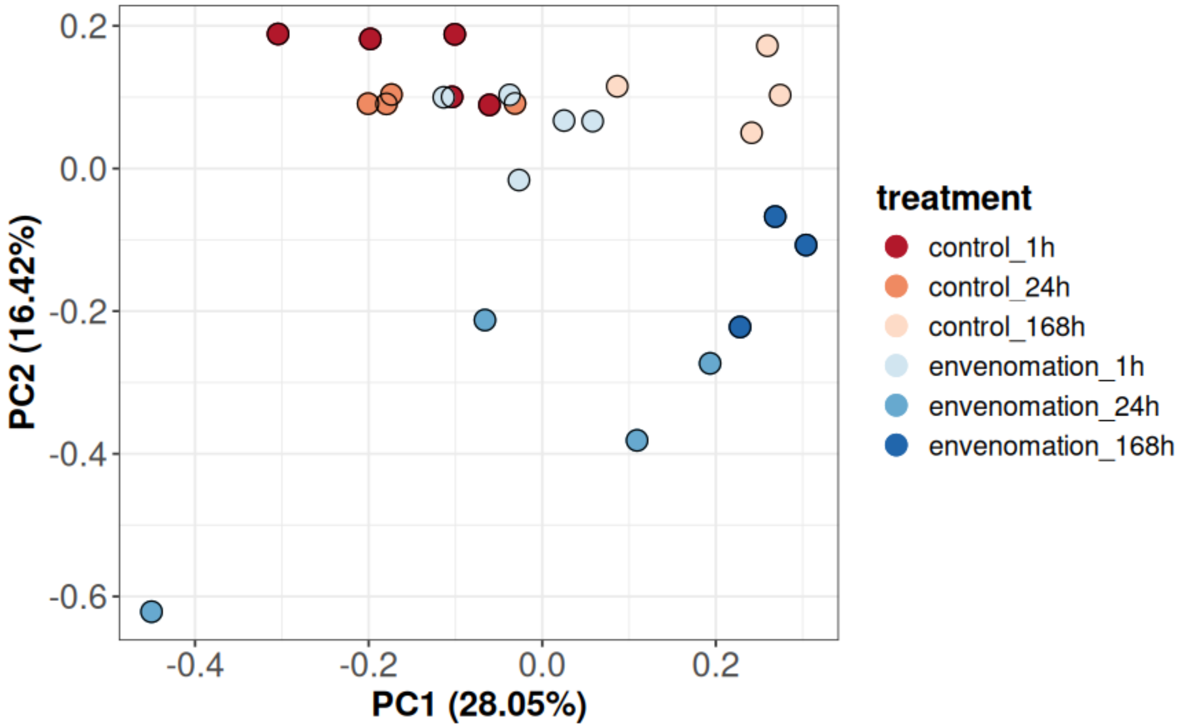


**Supplementary Fig. 1:** Principal component analysis (PCA) for identified venom sac proteins (n= 2031) based on label-free quantification (LFQ) intensity values showing the first two components (PC1, x-axis) explaining 28.05% and (PC2, y-axis) 16.42% of the variance within the dataset, respectively. Each treatment at each time-point is indicated by an individual colour.

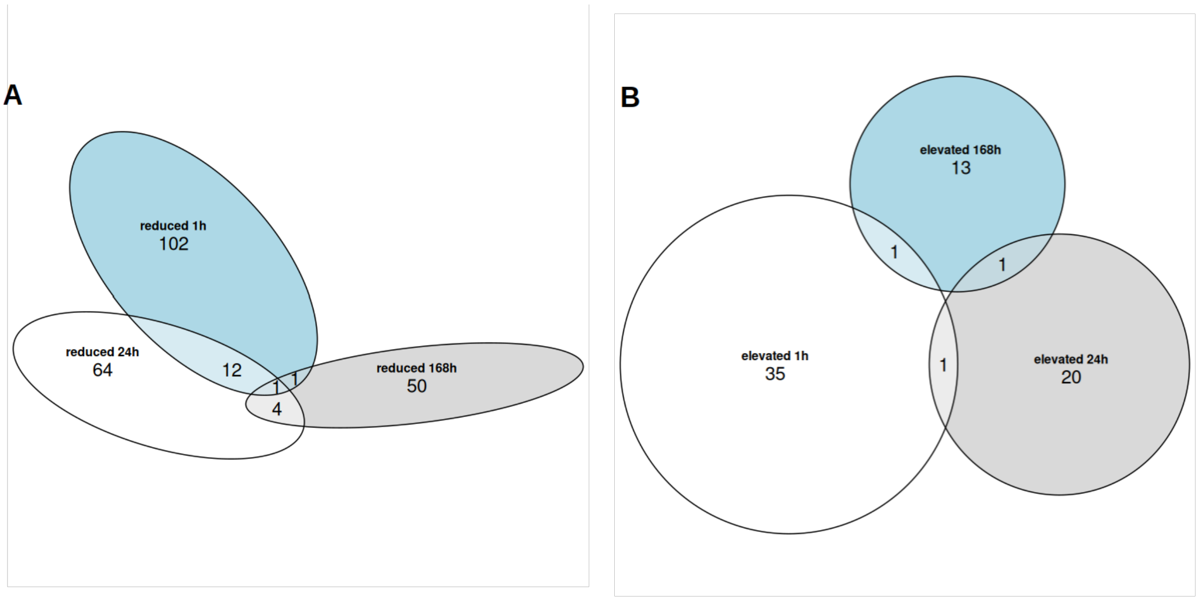


**Supplementary Fig. 2:** Overlap of significantly reduced or elevated venom sac proteins of *Bombus terrestris* workers post-envenomation across all time-points. **(A)** Reduced differently abundant venom sac proteins at all time-points. **(B)** Elevated differently abundant venom sac proteins at all time-points.
